# Global Genotype by Environment Prediction Competition Reveals That Diverse Modeling Strategies Can Deliver Satisfactory Maize Yield Estimates

**DOI:** 10.1101/2024.09.13.612969

**Authors:** Jacob D. Washburn, José Ignacio Varela, Alencar Xavier, Qiuyue Chen, David Ertl, Joseph L. Gage, James B. Holland, Dayane Cristina Lima, Maria Cinta Romay, Marco Lopez-Cruz, Gustavo de los Campos, Wesley Barber, Cristiano Zimmer, Ignacio Trucillo Silva, Fabiani Rocha, Renaud Rincent, Baber Ali, Haixiao Hu, Daniel E Runcie, Kirill Gusev, Andrei Slabodkin, Phillip Bax, Julie Aubert, Hugo Gangloff, Tristan Mary-Huard, Theodore Vanrenterghem, Carles Quesada-Traver, Steven Yates, Daniel Ariza-Suárez, Argeo Ulrich, Michele Wyler, Daniel R. Kick, Emily S. Bellis, Jason L. Causey, Emilio Soriano Chavez, Yixing Wang, Ved Piyush, Gayara D. Fernando, Robert K Hu, Rachit Kumar, Annan J. Timon, Rasika Venkatesh, Kenia Segura Abá, Huan Chen, Thilanka Ranaweera, Shin-Han Shiu, Peiran Wang, Max J. Gordon, B K. Amos, Sebastiano Busato, Daniel Perondi, Abhishek Gogna, Dennis Psaroudakis, C. P. James Chen, Hawlader A. Al-Mamun, Monica F. Danilevicz, Shriprabha R. Upadhyaya, David Edwards, Natalia de Leon

## Abstract

Predicting phenotypes from a combination of genetic and environmental factors is a grand challenge of modern biology. Slight improvements in this area have the potential to save lives, improve food and fuel security, permit better care of the planet, and create other positive outcomes. In 2022 and 2023 the first open-to-the-public Genomes to Fields (G2F) initiative Genotype by Environment (GxE) prediction competition was held using a large dataset including genomic variation, phenotype and weather measurements and field management notes, gathered by the project over nine years. The competition attracted registrants from around the world with representation from academic, government, industry, and non-profit institutions as well as unaffiliated. These participants came from diverse disciplines include plant science, animal science, breeding, statistics, computational biology and others. Some participants had no formal genetics or plant-related training, and some were just beginning their graduate education. The teams applied varied methods and strategies, providing a wealth of modeling knowledge based on a common dataset. The winner’s strategy involved two models combining machine learning and traditional breeding tools: one model emphasized environment using features extracted by Random Forest, Ridge Regression and Least-squares, and one focused on genetics. Other high-performing teams’ methods included quantitative genetics, classical machine learning/deep learning, mechanistic models, and model ensembles. The dataset factors used, such as genetics; weather; and management data, were also diverse, demonstrating that no single model or strategy is far superior to all others within the context of this competition.

## Introduction

Phenotype prediction is a grand challenge of twenty-first century biology (Azodi et al. 2019; Martinez 2023; National Research Council (US) 2010; U.S. National Science Foundation 2023). The ultimate goal of plant and animal breeding is to develop better phenotypes. Historically, breeding has been based on observation of phenotypes, recombining elite genetics and then selecting for superior phenotypes. With the advent of molecular genetic marker technologies and DNA sequencing, phenotypic selection has been augmented with genomic prediction and selection. In an agricultural context, genomic prediction (GP) and genomic selection (GS) have fundamentally altered plant and animal breeding by reducing generation interval, predicting traits that are too difficult and/or expensive to measure at population scale, and in the case of animals, improving the selection of sex limited traits (Bhat et al. 2016; Budhlakoti et al. 2022; Crossa et al. 2017; Desta and Ortiz 2014; Heffner et al. 2009; Johnsson 2023; Lorenz et al. 2011; Meuwissen et al. 2001; Washburn et al. 2020; Wiggans et al. 2017). Similarly, prediction models based on environmental and/or agronomic management factors have been critical tools for crop risk assessment, fertilizer and irrigation prescription, sustainability forecasting, climate change modeling, and other basic and applied research and decision making (Archontoulis et al. 2014; Challinor et al. 2018; Di Paola et al. 2016; Hammer et al. 2019; Hammer et al. 2002; Jones et al. 2003; Keating et al. 2003; Schauberger et al. 2017).

The terminology and importance placed on different factors can vary across applications. For example, plant and animal breeding tend to focus on the influence of genetic (G) factors on phenotypes, while sometimes including environmental (E) factors, and genotype-by-environment (GxE) interactions in their models. These fields take a plant-centric viewpoint where agronomic management (M) is included within E. Disciplines like agronomy and plant physiology on the other hand, often focus on M factors as distinct from E, taking a more farmer-centric approach where E is uncontrollable, and M includes all factors that can be manipulated by the researcher or producer. G may even be included as part of M in this framework, as the producer decides which cultivars to plant, in the same way that they determine fertilizer and irrigation rates.

Historically, applied agronomic prediction methods have focused on G, E, and/or M factors in relative isolation, but the past decade has seen a renewed interest in, and recognition of the need for, models that incorporate G, E, and M factors within a unified framework (Cooper et al. 2022; Guo and Li 2023; Jarquín et al. 2014; Kick et al. 2023; Kick and Washburn 2023; Li et al. 2021; Lopez-Cruz et al. 2023; Messina et al. 2023; Messina et al. 2018; Technow et al. 2015; Washburn et al. 2021). These approaches are sometimes referred to as GxE or GxExM depending on the context, but in practice they rely on diverse combinations of factors modeled in interacting and non-interacting ways.

In breeding, GP and GS approaches are well-suited for complex traits such as yield, which tend to have small single genetic variant (and per gene) effect sizes. They rely on genetic variation across the genome, as opposed to marker assisted selection approaches that use only a few selected loci known to be predictive of important variation for the trait of interest (Haley and Visscher 1998; Heffner et al. 2009; Meuwissen et al. 2001). Although many methods have been developed over the years to improve GP/GS accuracy, common linear and non-linear genomic prediction models, including Genomic Best Linear Unbiased Prediction (BLUP), Bayesian algorithms, Ridge Regression, Artificial Neural Networks and Decision Tree-based algorithms have often been shown to perform similarly across different species and trait combinations in cases where only G data is used, but differences can be large in some cases (Azodi et al. 2019; Charmet et al. 2020; Montesinos-López et al. 2021). Many genomic approaches have been developed to predict non-additive genetic effects, including modeling additive and dominant effects of individual markers, using multiple kernels (e.g. additive and dominance kernels) (Vitezica et al. 2013), and non-linear kernel regression methods (e.g. RKHS) (Morota and Gianola 2014). However, linear models often perform similarly to, or even outperform, non-linear algorithms, particularly when the trait has a predominantly additive genetic basis (Azodi et al. 2019; Montesinos-López et al. 2021).

When environmental variation is included in the model, linear, non-linear, and combined approaches that explicitly model environmental factors either statistically or mechanistically have been shown to improve predictive accuracy, particularly under certain stressful environments (Diepenbrock et al. 2022; Jarquín et al. 2014; Li et al. 2021; Ly et al. 2017; Millet et al. 2019; Technow et al. 2015; Washburn et al. 2021). Due to the complexity of experimental design in plant breeding trials, other practical strategies for analyzing multi-environment data have been developed, such as fitting linear mixed models with a two-stage approach for accelerating the speed of computation (Möhring and Piepho 2009; Rogers et al. 2021). The inclusion of dominance effects for hybrids and GxE effects for environment-specific genomic prediction are also likely to increase accuracy (Rogers et al. 2021; Rogers and Holland 2022). Another way to analyze multi-environmental trials is to group environments into what are termed mega environments (Lin et al. 2021).

Significant efforts have been made in the area of machine learning models for agriculture. Random forest (RF) models have been used to predict and analyze complex traits in plants by leveraging the collective intelligence of multiple decision trees (Azodi et al. 2019). RF excels at handling high-dimensional data and capturing non-linear relationships between predictors and responses. These attributes make it more suitable for capturing non-additive genetic effects and potentially environmental variation. Additionally, RF can provide variable importance measures, enabling breeders to identify key genetic markers and prioritize traits for selection. This versatility and robustness have made RF a valuable go to tool in prediction (Montesinos López et al. 2022). RF models have outperformed other methods for some traits and in some contexts of GP (Charmet et al. 2020; González-Recio and Forni 2011; Montesinos López et al. 2022).

Boosting is another method that is highly effective and is becoming widely used. Extreme gradient boosting is a scalable tree boosting system, that has been recognized for its computational speed and accuracy across a range of prediction problems (Chen and Guestrin 2016). Gradient boosting of decision trees, for example XGBoost (Chen and Guestrin 2016) and LightGBM (Ke et al. 2017), and related RF methods have been observed to significantly outperform deep-learning models in several plant and non-plant tabular datasets and they typically require fewer computational resources (Borisov et al. 2022; Danilevicz et al. 2021; Gill et al. 2022; Grinsztajn et al. 2022; Shwartz-Ziv and Armon 2022).

Predicting phenotypes often requires combining multiple heterogeneous sources of information (Xu et al. 2019). This can complicate the selection of a meaningful subset of features to use as predictors. Deep learning models are highly effective in handling large and diverse input data and modeling complex nonlinear relationships, surpassing traditional modeling approaches (Khaki and Wang 2019). Additionally, they can incorporate specialized architectures such as long-short-term memory (LSTM), which is particularly suitable for time series data due to its capacity to capture and retain long-term dependencies (Malhotra et al. 2015; Shook et al. 2021). Deep learning models have shown mixed results when applied to GP scenarios (Kick et al. 2023; Montesinos-López et al. 2021; Washburn et al. 2021). Most successful applications of deep learning involve enormous datasets of a scale beyond those typically used in plant breeding and other agricultural scenarios, which has potentially limited the true potential of these methods in agricultural phenotype prediction. The computational requirements of these methods are also potentially limiting even with relatively small datasets, but the computer gaming industry has resulted in wider access to inexpensive graphical processing units (GPU) making these methods arguably less expensive than some traditional BLUP approaches that require prohibitively large amounts of memory. Other approaches have involved a combination of deterministic models, expert knowledge, and deep learning to model environmental stress in agriculture (Cvejoski et al. 2021).

Another important class of methods are ensembles of different model types. Ensembles allow the combination of other types of methods, including any of those discussed above, into a single prediction and can often outperform each method on its own (Kick and Washburn 2023; Shahhosseini et al. 2020). Ideally, ensembling allows for reduced error (Zhou 2015) and greater robustness by pooling the predictions of a diverse set of models. Ensemble learning has been used for purposes ranging from optimizing genetic transfer to genetic prediction, but substantially more research has focused on developing and comparing single model performance than ensemble approaches (Azodi et al. 2019; Hesami et al. 2020; Liang et al. 2021).

One significant challenge to improving phenotype prediction methods is the lack of publicly available datasets containing G, E, and M factors with which to experiment and develop new prediction approaches. In 2013, collaborators from universities, government, farmer associations, and industry recognized the need for this type of data and formed the Genomes to Fields Initiative (G2F) Maize Genomes by Environment (GxE) project centered around maize yield trial data. To date, this project has evaluated over 180,000 unique plots, 5,000 maize hybrids, and 280 environments (Lima et al. 2023a; Lima et al. 2023b; McFarland et al. 2020). Phenotypic, genetic, environmental, and management data have been collected following standard protocols and uploaded annually to a joint repository for public use. The individual datasets have resulted in many publications and advancements in our knowledge and understanding of maize genetics, phenotypes, and environmental responses (Bai et al. 2019; DeChant et al. 2017; Falcon et al. 2020; Gage et al. 2017; Gage et al. 2019; Lopez-Cruz et al. 2023; Morales et al. 2020; Rogers et al. 2021; Sekhon et al. 2020; Stewart et al. 2019; Wiesner-Hanks et al. 2018; Wiesner-Hanks et al. 2019; Wu et al. 2019). The datasets have also been used individually for multiple studies on phenotype prediction (Anche et al. 2020; Anderson Ii et al. 2019; Jarquin et al. 2021; Kick et al. 2023; Kick and Washburn 2023; Rogers and Holland 2022; Washburn et al. 2021; Westhues et al. 2021; Westhues et al. 2022; Winn et al. 2023).

While the G2F dataset has been shared with the public throughout the project, and continues to be updated as new data is collected and processed, barriers to using the data have persisted (Lopez-Cruz et al. 2023). These barriers included the vast size and complexity of the dataset, the many different data types collected, each of which requires some unique domain knowledge to interpret, and the fact that many potential users were simply unaware of the data. Additionally, to make the dataset useful for a wide range of studies, curation has been kept to a minimum to allow each researcher to determine what data is most valid for their specific use case.

One of the original goals of the G2F GxE project had been to use the data for the development of phenotype prediction methods. From November 15, 2022 to January 15, 2023 the G2F GxE hosted its first ever yield prediction competition with the objectives of expanding the number of people and domains working with the data set and stimulating the development of new and innovative phenotype prediction models and strategies. Competition challenges are very common in some domains of science (e.g., computer science, data science, etc.), but they are relatively uncommon in plant science, breeding, genetics, and related disciplines. For the G2F community, this was a completely new endeavor. Planning and advertising for the competition began more than a year in advance, along with an extensive effort to format and curate the existing G2F data to date into a form that would be more accessible to competition participants and future researchers. The goal for competition participants was to use the G2F GxE data from 2014 to 2021 (termed the training set) to predict the grain yield results from the G2F GxE 2022 season (termed the testing set) which had not yet been publicly released. This represented one of the most difficult prediction scenarios in crop agriculture because it required prediction on many new genotypes (hybrids not included in the training set) in new environments (a new year not included in the training set). Of the hybrids for prediction in 2022, only 8% had been previously tested in the G2F GxE project. Many of the field locations had been used previously, but no yield performance data at any of the locations for 2022 was provided to the participants.

This manuscript describes the results of the competition, including the different modeling strategies employed and their effectiveness. To encourage industry participation, teams were allowed to keep their methods confidential (see Materials and Methods for details). Teams were also allowed to publish their models on their own rather than being included in this joint manuscript. Several teams elected to do so and their manuscripts are in various stages of preparation, submission, and publication along with non-participants using the data independently (Fernandes et al. 2024; Ge et al. 2024; Khalilzadeh et al. 2024; Lopez-Cruz et al. 2024). This manuscript includes the results from all eligible participant teams as well as detailed methods, results, and code for 17 of the participant teams, including a majority of the top ten and others distributed across the range of scores, who opted to participate. Particular emphasis is given to the winning team’s approach, methods, and results.

## Materials and Methods

### Competition datasets

A substantial curation, quality control, formatting, and standardization effort on the G2F data was carried out to lower the barrier to entry and facilitate the participation of individuals from as many different disciplines and backgrounds as possible. Compilation of the phenotypic data set began by combining the “clean” versions of the raw trait data files posted as DOIs for public distribution. The “clean” trait data files had undergone automated filtering to remove extreme outliers, as described in the DOI readme files and dataset publications (Lima et al. 2023a; Lima et al. 2023b; Lima et al. 2023c). Trait data only from the “main” Genomes to Fields experiments were included; data from several smaller “side” experiments were removed. Sub-experiment and field block information was recovered for missing information whenever possible. Trait and meta-information column names were harmonized among years. Hybrid names were harmonized between years of the training data and between training and testing sets. Duplicate records were dropped.

Metadata files were curated in a similar way as the phenotypic data; files were downloaded from DOIs and compiled. Environment names were harmonized among the years, and between metadata and phenotypic data to ensure consistency. Only those environments present in the phenotypic data were kept in the metadata. Any specific information related to a particular environment was added to the "Comments" column. Columns for "City", "Farm", and "Previous Crop" were also harmonized, double-checked and updated, when possible. Soil data collection only began in 2015. For all years after that, files were obtained from the DOI and compiled. As with the metadata, environment names were standardized across all years and between phenotypic data, and only environments present in the phenotypic data were included in the soil data file. For some environments, data was unavailable in the DOI as the results were returned after the DOI release. In such cases, the data was retrieved from the laboratory responsible for the analysis and included in the competition DOI release (Lima et al. 2023c).

For the genotype data, the maize Practical Haplotype Graph (PHG) was used for variant calling (Bradbury et al. 2022). The PHG aligns sequencing reads with genome assembly sequences to impute genotypes based on stored haplotypes. The Maize 2.1 PHG haplotypes came from 86 genome assemblies which were aligned to the B73 v5 assembly (Hufford et al. 2021) with anchorwave (Song et al. 2022). The assemblies came from MaizeGDB and other sources (Bornowski et al. 2021; Woodhouse et al. 2021; Yang et al. 2019). The B73 genome was divided into nodes (both genic and intergenic) known as reference ranges, using the annotations from Zm-B73-REFERENCE-NAM-5.0_Zm00001eb.1.gff3. The ends of reference ranges were selected as regions with 10 or more conserved base pairs in 23 or more of the 25 NAM genome assemblies (Hufford et al. 2021). Genic haplotypes having 0.0001 or lower divergence and intergenic haplotypes having 0.001 or lower divergence were collapsed into consensus haplotypes.

Due to changing technologies over the years, the inbred sequence data (which is combined to create the hybrid genotypes) collected for the G2F project has come from different methods in different years of the project. The 2014-2017 genotyping was done using a GBS (genotyping by sequencing) protocol (Elshire et al. 2011) with the ApeKI restriction enzyme. The 2018-2019 genotyping was performed with ∼5x coverage skim sequence. The 2020-2021 genotyping used Exome capture in combination with GBS controls using ApeKI. The 2022-2023 genotyping used GBS with PstI-MspI. To create the genotype calls from the G2F data, inbred reads were aligned to the PHG to identify haplotypes matching each read. The haplotype path through the graph was then imputed and used to identify variants. Only positions contained within the 600 k SNP genotyping array (Unterseer et al. 2014) were considered. CrossMap was used to uplift array positions to v5 coordinates and positions were dropped if they could not be uplifted, were missing in 21 or more assemblies or were monomorphic in all assemblies (Zhao et al. 2014). The final set contained 437,214 variant positions. Hybrid names in the genotypic data set were harmonized with the hybrid names in training and testing trait data sets.

Weather data was downloaded from the NASA Power website (https://power.larc.nasa.gov/) for the locations and years in the training and testing sets. An estimate was made for locations where the exact GPS field coordinates are unknown. Competitors could also use other sources of weather data, including the weather data available in the DOI of each year of the project; each environment is equipped with a weather station (WatchDog 2700 Weather Station) that records various weather parameters every 30 minutes during the growing season, including air temperature, humidity, solar radiation, rainfall, wind speed and direction, soil temperature, and soil moisture. Environmental covariate data was also given to the participants. Details of how these environmental covariates were generated are described in (Lopez-Cruz et al. 2023). It is important to note that these covariates were defined at the year-location level; therefore, these did not vary within year-location.

The final curated dataset used in the competition, including the observed phenotypic (test set) values that were not released until after the competition, is publicly available for further use (Genomes to Fields 2023; Lima et al. 2023c). The released dataset also contains an extensive readme file with details about each component of the dataset exactly as they were presented to the competition participants. The training and testing set data and a short numerical summary of the contents of each can be found in Table 1.

**Table 1:**
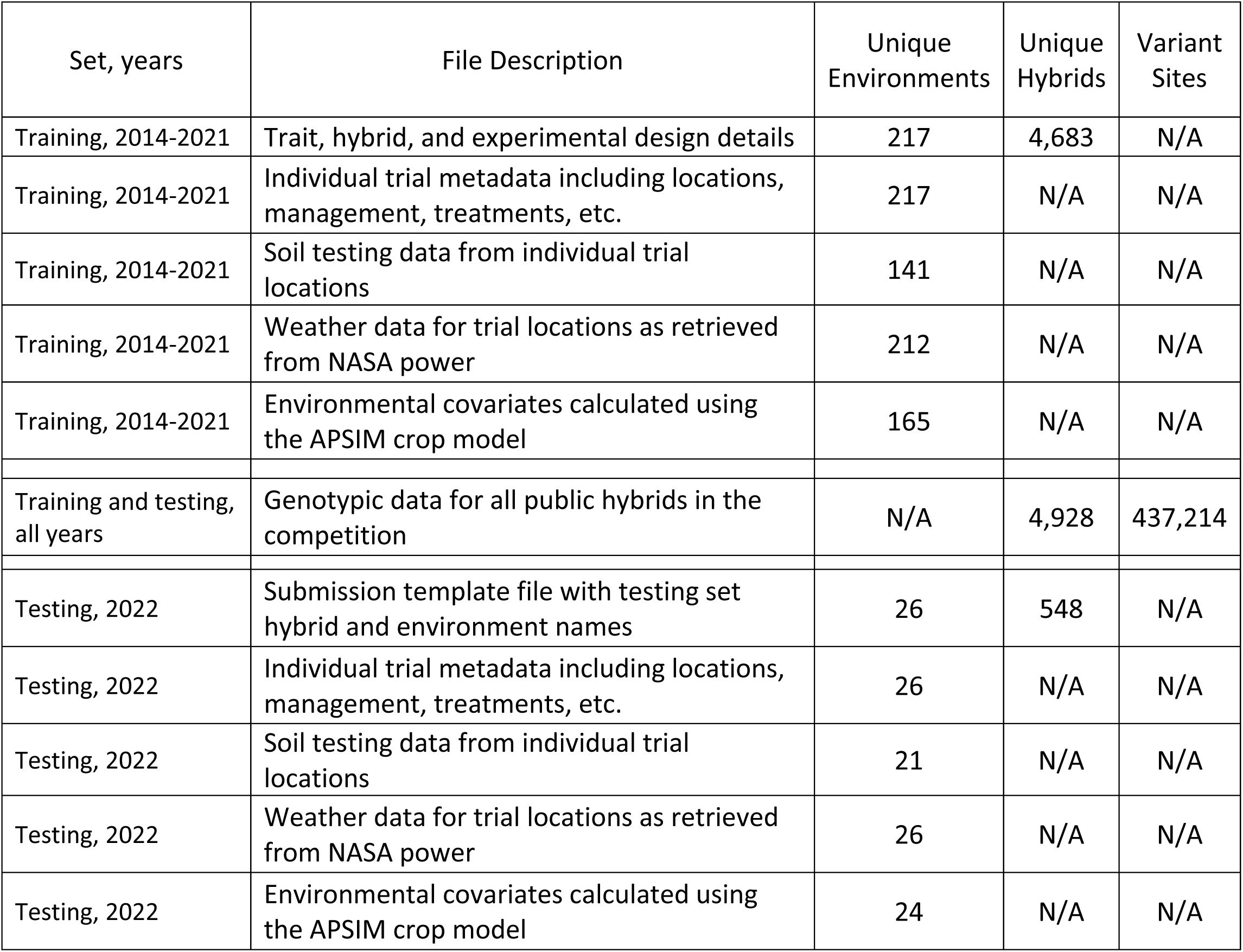
Data files made available for the competition.

### Competition procedures and methods

The competition hosted a website for advertising, information, and competition rules and utilized the EvalAI platform (Yadav et al. 2019) for submissions, evaluations, and leaderboard hosting (see File S1 which contains a preserved version of all competition website information). EvalAI is an open-source platform for evaluating and comparing machine learning and artificial intelligence outputs and algorithms. The platform allows the host to control the number of challenge phases, dataset splits and leaderboard visibility by cloning a git repository that contains the configuration files in the YAML language. The competition rules were described in detail in the website materials and the EvalAI challenge webpage. Individuals and teams from any institution were encouraged to participate but could only be part of one team in the competition. Any model or strategy was permissible as long as it relied only on data provided by the competition or external data that was publicly available by February 1, 2022. This date was chosen to prevent competitors from using private data or data collected during the 2022 growing season, which is the season they were trying to predict. The submitted predictions were to be of absolute grain yield for each hybrid for each test environment. These were reported in megagrams (metric tonnes) per hectare (Mg/ha) with the standard 15.5% moisture adjustment used for maize in the U.S. The evaluation metric used was the average of the root mean squared error (RMSE) calculated for each environment. In other words, for each submission the RMSE for each environment was calculated individually and then these RMSE values were averaged for a final score. The winner was the team with the lowest average RMSE value. To be eligible to win the competition and receive the cash prize, participants were required to commit to publishing their model code and results after the competition (on their own or as part of this combined manuscript). To allow greater participation from industry groups, an option was provided for participation in the competition without publication of methods, but these groups would be ineligible for the prize and winning title.

The competition began on November 15, 2022 at which time the training data, testing data, and submission template (Table 1) were provided to all teams. Each team was allowed to make five total submissions through the EvalAi system by the close of the competition on January 15, 2023. Results (Mean RMSE) were evaluated automatically using a python script hosted in the EvalAI remote servers and displayed on the competition leaderboard within seconds of submission. The calculation of RMSE was performed within each environment and then averaged across all environments. The EvalAI leaderboard provided participants with feedback on their submissions that could be used to improve their model and also allowed for greater interaction between participants. At the end of the competition, the top three teams on the leaderboard were contacted and required to provide their code to the organizers for a complete evaluation. This process ensured that the winning team was following competition rules as outlined, and that their results could be re-created in the hands of the organizers. There was one team (DataJanitors) made up of a few of the organizers of the competition, they were allowed to make submissions “for fun” as long as they followed all the normal competition rules (not using the truth values), but they would not and were not considered part of the competition or eligible to win. However, their methods and results are included here.

### Individual team methods

The prediction methods used by different teams in the competition were extremely diverse. Methods ranged from traditional BLUP and linear mixed model approaches, to deterministic models, to classical machine learning methods, random forest models, ridge regression, gradient boosted decision trees, and various deep learning neural network approaches (Abadi et al. 2016; Bradbury et al. 2007; Breiman 2001; Butler et al. 2017; Chen and Guestrin 2016; Chollet 2015; Jarquín et al. 2014; Ke et al. 2017; Paszke et al. 2019; Pedregosa et al. 2011; Pérez and de los Campos 2014; Wright and Ziegler 2017). Many of the methods were also applied jointly using ensemble approaches. Most methods were implemented using packages in Python and R (R Core Team 2021; Van Rossum and Drake 2009). Some teams used feature selection methods, GWAS, and/or external datasets with no direct relationship to the G2F data in attempts to improve their model’s predictive accuracy. The methods and approaches used by each team are described in detail in Files S2 and links are provided there to the code developed by each participating team.

### Winning team methods

The most accurate prediction model in the competition was submitted by the Corteva Latin America Corn (CLAC) team. It consisted of averaging two model outputs, namely A and B. Model A was a univariate linear mixed-effects model, where the fixed effects included location meta-data: location (state, station), irrigation, treatment, and previous crop. No year or year-location effects were accounted for. The random effect consisted of a polygenic genetic term, with a relationship matrix calculated as an arccosine kernel. Predictions from model A were generated from the fixed effects and genomic values estimated for the 2022 data.

Model B consisted of a location specific model using environmental variables and the location meta-data to predict the environmental means, while relying on an index from a multivariate GBLUP to predict the genetic component. The environment means model fitted the mean yield of environments as a function of environmental variables and the location meta-data, fitted in two steps: First, it estimated biased composite predictor as the average of three sub-models: 1) a random forest of the environmental factors, 2) a ridge regression of environmental factors, 3) a least-squares of the location meta-data; Second, it computed the unbiased estimator, using a linear regression to remove the shrinkage from the composite prediction. Model B’s predicted genomic values were inferred from a selection index. The predictions started from fitting an unstructured model of the observed environments as

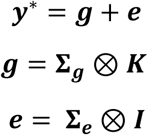

where **y\*** were standardized and spatially adjusted phenotypic values; **g** and ***e*** were the genetic effect and residuals of each corresponding phenotype, respectively. The genetic covariance matrix **Σ_g_** contained the variances of each environment in the diagonal and covariances between pair of environments in the off diagonal. The residual covariance **Σ*_e_*** was a diagonal matrix containing the residual variance for each environment. K was the genomic relationship matrix, where the pairwise relationship among individuals was calculated from an arc-cosine function using genomic information (Cuevas et al. 2019). The output of the model consisted of predictions of every individual in every environment. The predicted genetic merit for the k^th^ 2022 location (u_k_) was estimated as a linear combination of the genomic values of the observed locations as:

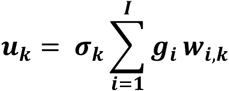

where the scalar w_i,k_ corresponded to the weight of the i^th^ location to predict the k^th^ location of 2022, and σ_k_ was the predicted standard deviation for the k^th^ location. The genotypic standard deviation of 2022 locations were predicted from the phenotypic standard deviation of location as a function of environmental covariates using random forest (*σ* = *RF*(*W*) + *e*). The weights (w_i,k_) were based on the deterministic accuracy of i^th^ predicting k^th^ location and the geographical location, such that locations in the same state and station would have higher weights than location further away, and locations with individuals more related to those in the k^th^ location will also have higher weights.

All computations were done in R. Linear mixed models were fitted using the R package bWGR 2.1 (Xavier 2019; Xavier and Habier 2022; Xavier et al. 2020). The univariate model was described by Xavier (2019) and the multivariate model by Xavier and Habier (2022). The random forest used the R package ranger (Wright and Ziegler 2017).

## Results and Discussion

### Competition participation

Overall, the competition had 241 registrants comprising 128 teams from at least 19 countries and every inhabited continent (Figure 1A). Registrants came from academic institutions, government, non-profits, and private companies (including major seed companies, small startups, biotechnology companies, and international tech giants, Figure 1B). The mean number of members of a team was 1.9 with most teams consisting of only one individual and the largest team consisting of 10. Other demographic information on the participants was not recorded. By the end of the competition, 30 teams (excluding a few test teams and erroneous submission) successfully submitted a model on the leader board. That represents a 77% drop out rate, which was congruent with expectations given the significant effort and time required to develop a successful model with a large and heterogeneous data set of this kind. Interestingly, while 62% of the original registered teams contained only one member, 70% of the teams with successful submissions had two or more members and the mean number of members in these final teams was 2.9.

**Figure 1.**
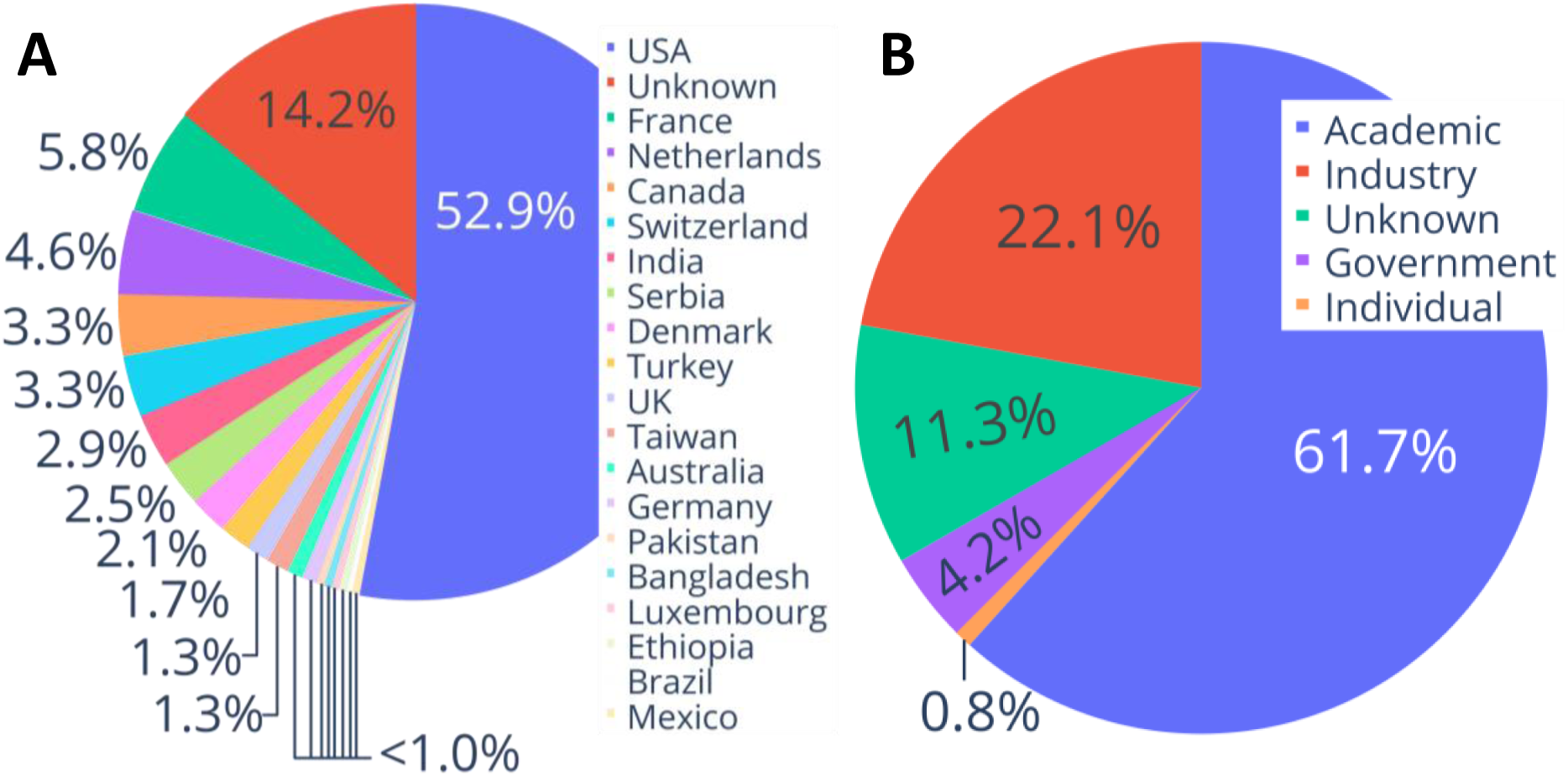
Competition Participation by: A) Country, and B) Institution type.

### Model accuracies

One important difference between this competition and other model development studies is the fact that participants did not have access to the observed values. The best practice is for researchers to have a “validation set” in addition to training and testing sets, and iteratively improve their models on that set, saving the testing set for final testing and evaluation only. Ideally, the number of times a model is challenged with the testing set (and potentially changed by the researcher to perform better) should be minimized. However, this is difficult to implement for even the most well-meaning researchers due to small sample sizes, concerns about representation in sets, and the fact that if one’s model performs very poorly on the testing set, it may be unpublishable or indicative of improvement areas. In this competition, participants had up to five chances to challenge their model against the testing set; four of these could be used as feedback to improve the model for the fifth attempt. The motivation for this is described in the methods section. While this allowed for some improvement based on testing set feedback, the number of possible iterations was much lower than what one could do with testing set access and the feedback given was minimal (a single accuracy score).

Of the teams that submitted models, 67% made all five submissions, 10% made four, 7% made three, 3% made two, and 13% made only one. There was a significant (p=0.024) Spearman correlation of -0.412 between the final rankings of teams and the number of submissions they made, indicating that teams with more submissions did better on average than teams with fewer (the small sample size of 30 teams should be considered in these interpretations). Even so, it did not appear that having five submissions played a major role in model improvements. Most submissions were made near the end of the competition (see Figure S1) with 71% of submissions in the final week and almost 60% in the last three days. When submissions from each team were compared to their previous submission (e.g., submission 5 vs. submission 4, 4 vs. 3, etc.), 52% of submissions performed better than the previous submission and 48% performed worse. When only the first and final per team submissions were considered, 46% of final submissions resulted in improvement while 54% resulted in lower accuracy. The reasons behind this are not clear but it may have been that more committed teams simply made more submission attempts. The winning team was in first-place for most of the competition with only a few days when the second-place team was in the lead.

The density distribution of the final model accuracy scores from the competition can be seen in Figure 2. Only the top 10 models are highlighted in the figure with a red “x”. The top 10 best scoring models are relatively close together: within 0.215 Mg/ha. As a percentage of the average yield of the testing set, this difference in model errors is 2.13%, indicating a small improvement from the 10^th^ place model to the 1^st^ place, but one that could still be considered useful for many applications. Looking across the competition (with the exclusion of one outlier model), the 1^st^ ranked team’s score represents a 9.42% improvement.

**Figure 2.**
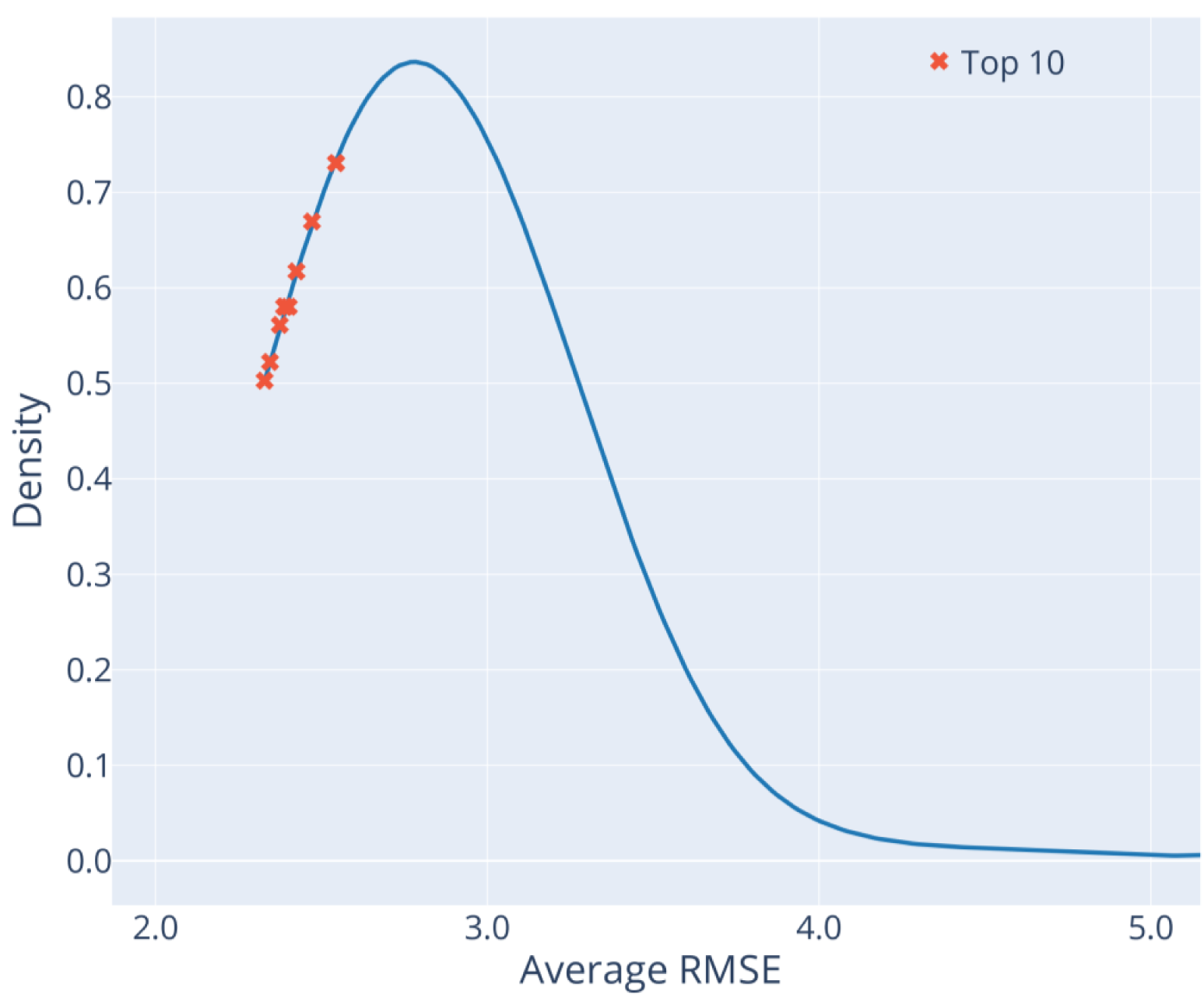
Density distribution of final competition scores. The top ten best scores are shown with an “x”. The figure is truncated around 5.0 Average RMSE for simplicity.

While many measures of model accuracy could have been used, the average RMSE metric was chosen for determining the competition winner due to its simplicity and common use in modeling, machine learning, and deep learning (from which the organizers hoped to attract participants). The best evaluation metric in any study depends on the goals and desired applications. While RMSE, MSE, and similar metrics are more widely used the broader field of modeling, Pearson correlation coefficient (r) is generally preferred in breeding since knowing the best or worst performing lines is the application. No metric is perfect, Pearson r allows for wildly different absolute values for example (in fact one submission with absolute values far outside of reasonable corn yields would have scored much better if Pearson r were the metric, see Table S1). The use of Pearson r is recommended for future competitions, if the goal is more breeding focused.

To better explore the competition results, numerous metrics were calculated and compared (see Tables S1 and S2). The scores and rankings based on the average RMSE and other common metrics for the top 20 teams in the competition are shown in Table 2. Rankings that differed from the average RMSE ranking are shaded. Although the absolute ranking of teams changed considerably depending on the metric used, the top and bottom few models were relatively consistent across metrics, with mostly small rank differences observed. The best average RMSE model (the competition winner) remained the winner when evaluated with most other common metrics although it placed 2^nd^ for Global Pearson Coefficient (r), a metric that considers accuracy across all environments jointly rather than separately. While the top and bottom teams were fairly robust across metrics, the rank differences seen, which are in some cases large (for example moving from 15^th^ place to 4^th^ place) illustrate the importance of carefully and deliberately choosing metrics when comparing model accuracies.

**Table 2:** Ranking of the top-20 teams.

| Team Name | Average of within-environment scores |  |  |  | Across-environment scores |  |  |  |
| --- | --- | --- | --- | --- | --- | --- | --- | --- |
|  | RMSE |  | Pearson r |  | RMSE |  | Pearson r |  |
|  | Score | Rank | Score | Rank | Score | Rank | Score | Rank |
| CLAC | 2.329 | 1 | 0.357 | 1 | 2.458 | 1 | 0.631 | 2 |
| igorkf | 2.345 | 2 | NaN | NaN | 2.464 | 2 | 0.600 | 6 |
| phenomaize | 2.374 | 3 | 0.238 | 8 | 2.470 | 3 | 0.617 | 4 |
| UCD_MegaLMM | 2.387 | 4 | 0.338 | 3 | 2.505 | 6 | 0.616 | 5 |
| CGM | 2.391 | 5 | 0.353 | 2 | 2.490 | 5 | 0.587 | 7 |
| Purdue | 2.402 | 6 | 0.161 | 15 | 2.488 | 4 | 0.631 | 3 |
| SmAL | 2.425 | 7 | 0.146 | 17 | 2.525 | 7 | 0.586 | 8 |
| ML_APT | 2.472 | 8 | 0.191 | 13 | 2.600 | 8 | 0.564 | 10 |
| MPB_Group | 2.544 | 9 | 0.255 | 6 | 2.741 | 11 | 0.494 | 13 |
| AlBreeding | 2.544 | 10 | 0.220 | 9 | 2.758 | 14 | 0.439 | 15 |
| AgroStat | 2.562 | 11 | 0.100 | 20 | 2.646 | 9 | 0.554 | 11 |
| arulrich | 2.575 | 12 | 0.205 | 12 | 2.726 | 10 | 0.510 | 12 |
| DataJanitors | 2.587 | 13 | 0.256 | 5 | 2.752 | 12 | 0.644 | 1 |
| CropsAreCool | 2.616 | 14 | 0.161 | 14 | 2.805 | 15 | 0.413 | 19 |
| AllModelsAreWrong | 2.646 | 15 | 0.272 | 4 | 2.754 | 13 | 0.575 | 9 |
| agAdaptAR | 2.685 | 16 | 0.151 | 16 | 2.848 | 18 | 0.400 | 22 |
| CropEnthusiast | 2.710 | 17 | 0.116 | 19 | 2.844 | 17 | 0.437 | 16 |
| DeepCropVision | 2.739 | 18 | -0.131 | 29 | 2.829 | 16 | 0.388 | 23 |
| supermanwasd | 2.746 | 19 | 0.243 | 7 | 2.915 | 21 | 0.409 | 21 |
| BioSense | 2.747 | 20 | 0.131 | 18 | 2.892 | 19 | 0.410 | 20 |

Shading = Rankings that differ from the average RMSE ranking

Simple average-based ensemble models were also created and tested using combinations of submissions. A model based on all valid submissions combined produced an RMSE of 2.580 and would rank 13^th^ within the leader board. A model using the best submission from each of the top 12 teams performed better than the best submission in the competition. This trend continued as the number of top teams decreased with the highest performing ensemble model being a combination of the top two teams with an average RMSE score of 2.288. However, the best performing ensemble based on the average of Pearson r environment scores was a model including the top 5 teams with a score of r = 0.369. In summary, some simple ensemble models marginally outperformed the top submissions in the competition.

Some participants focused on predicting environmental means, reasoning that getting those right would be more important than genetics, since the environments were so diverse and RMSE scores favor absolute values over genotypic rankings. The box plots in Figure 3A demonstrate that the predictions on average had narrower distributions than the observed values for each environment. Predicting environmental means proved challenging for all teams, particularly for certain environments (Figure 3B, Figure 4). For example, WIH1_2022 (Wisconsin) had the highest average yield of all the environments in the test set and was consistently underpredicted by nearly every team (Figure S2). Most midwestern environments were predicted to have similar average yields to one another. Two exceptions were the IAH2 (Iowa), and NEH1 (Nebraska) locations which performed worse than predicted. IAH2 also had the greatest observed variation of the locations. In contrast, several of the southern and northern locations, including NCH1 (North Carolina) and Texas and New York, were predicted very well on average, GAH1 (Georgia) being an exception. In summary, the most difficult environments to predict were those on the extremes with the highest and lowest observed yields and the team’s predictions tended toward the mean of all environments (Figure 3B).

**Figure 3.**
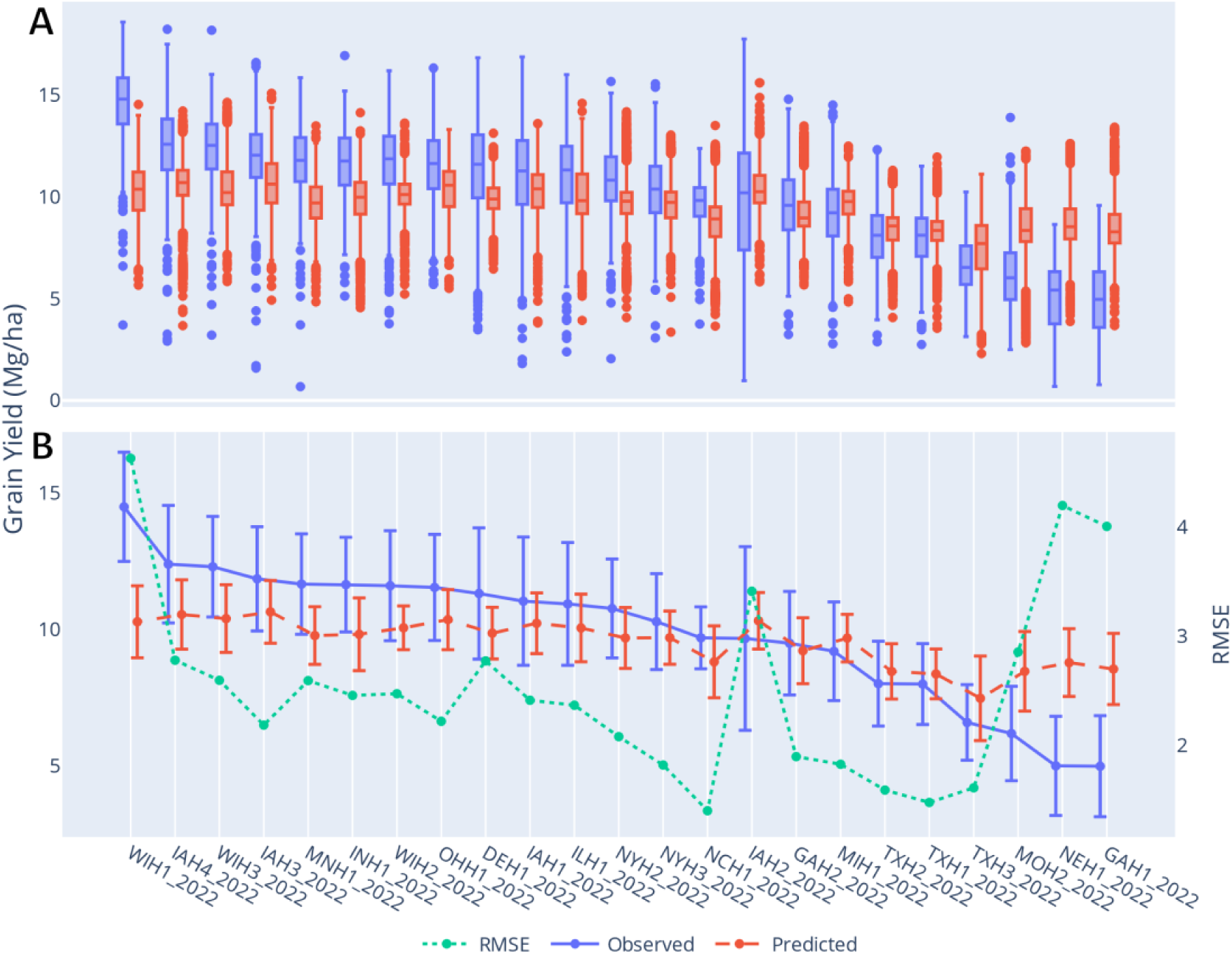
Test set environmental distributions based on the observed and predicted values from all teams (excluding the final team on the leaderboard due to significant outliers) shown as: A) Box plots, and B) Environment means with standard deviations as error bars and average RMSE scores.

**Figure 4.**
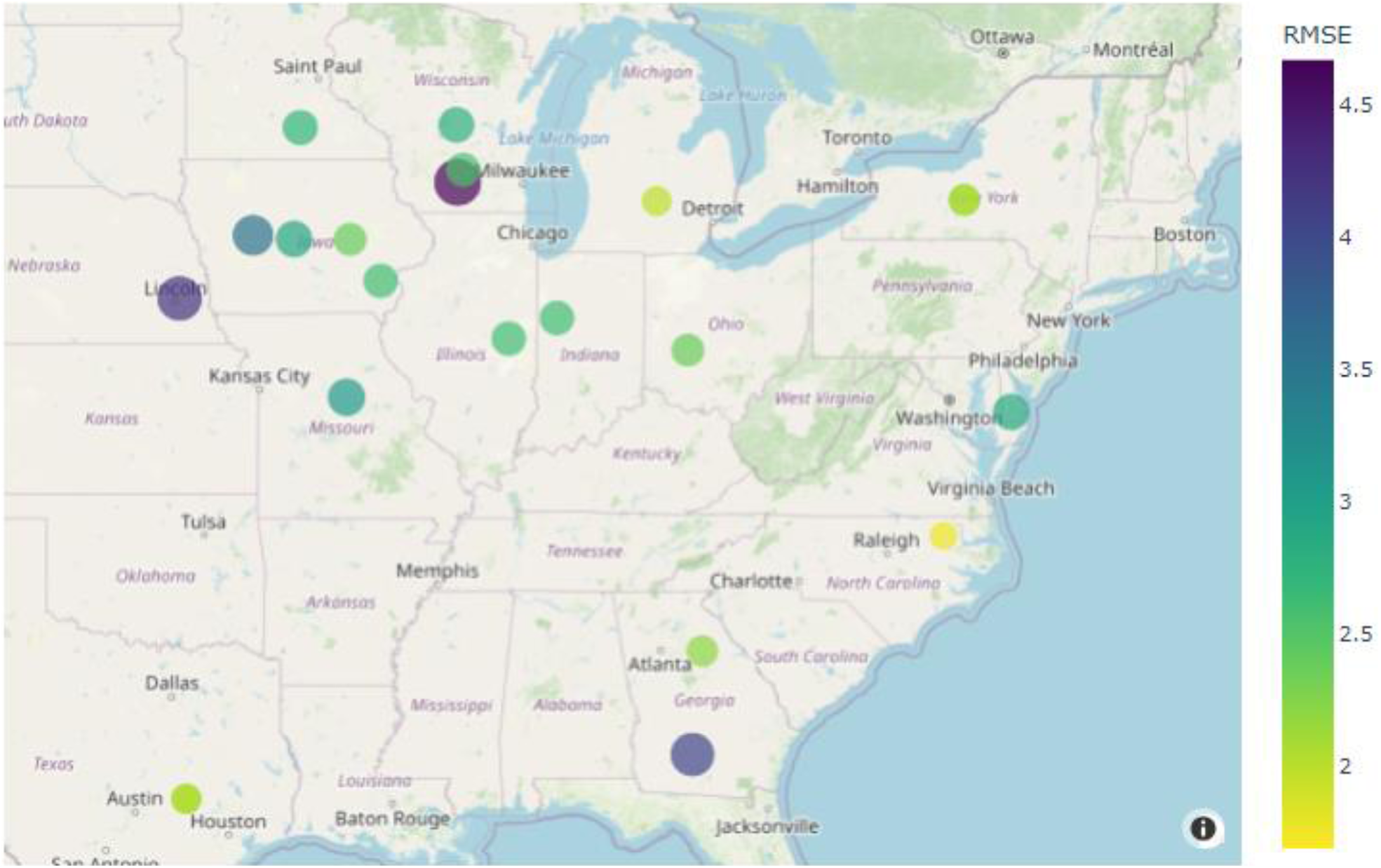
Average of all team’s (excluding the final team on the leaderboard due to significant outliers) per environment RMSE scores plotted on a U.S. map. Both the color and size of the dots represent RMSE scores.

### Modeling strategies

Diverse modeling strategies, including different ways of pre-processing the data, were applied by teams in the competition. These approaches included variations on traditional BLUPs and linear mixed models, deterministic models, classical machine learning, random forest, ridge regression, gradient boosting decision trees, deep learning neural networks, and many different combinations of these approaches (See File S2). Not every participating team chose to reveal the details of their model but 27 of the 30 teams responded to end-of-competition polls with details about the model types they used. The most highly used model types were within the category here described as classical machine learning, with 63% of respondents indicating they used modeling approaches within this category (See Table 3). Although this was the highest used model type, it was often used in conjunction with other models, and in some cases was primarily used in pre-processing steps with a different model type being used for the final prediction. Still, this indicates the important role that classical machine learning models play in modern phenotype prediction. For the subset of teams that chose to release their methods in greater detail within this manuscript, Random Forest, Gradient Boosting, and Ridge Regression, in that order, were the most commonly used classical machine learning methods (See Table S3).

**Table 3:** Modelling strategies used by team.

|  | Model Type |  |  |  |  |  |
| --- | --- | --- | --- | --- | --- | --- |
| Team Name | Rank | Classical<br>Machine<br>Learning | Linear/<br>Mixed/<br>BLUP | Deep<br>Learning/<br>Neural<br>Net | Other<br>Strategy | Ensemble<br>Model |
| CLAC | 1 | X | X |  |  | X |
| igorkf | 2 | X |  |  |  |  |
| phenomaize | 3 |  | X |  |  |  |
| UCD_MegaLMM | 4 |  | X |  |  |  |
| CGM | 5 |  | X |  |  |  |
| Purdue | 6 | X | X |  |  | X |
| SmAL | 7 |  |  | X | X |  |
| ML_APT | 8 | X | X |  |  | X |
| MPB_Group | 9 |  | X | X |  |  |
| AlBreeding | 10 | X | X | X |  | X |
| AgroStat | 11 | X |  |  |  | X |
| arulrich | 12 | X |  |  |  |  |
| DataJanitors | 13 | X | X |  |  | X |
| CropsAreCool | 14 | X |  |  |  | X |
| AllModelsAreWrong | 15 | X | X |  |  | X |
| agAdaptAR | 16 | X |  | X |  | X |
| DeepCropVision | 18 |  |  | X |  | X |
| supermanwasd | 19 |  |  |  |  | X |
| BioSense | 20 | X |  |  | X |  |
| Kernel of Truth | 21 | X | X |  |  | X |
| AlMaize | 24 | X | X | X |  | X |
| EnBiSys | 25 |  |  | X |  |  |
| Genetwister | 26 |  | X |  |  |  |
| gartybois | 27 | X | X |  |  |  |
| Niche Squad | 28 |  |  | X |  |  |
| uwaBioinfo | 29 | X |  |  |  | X |
| TinyAfrica | 30 | X |  | X |  |  |
| Percentage of Teams Using Strategy |  | 63% | 52% | 33% | 7% | 52% |

The second most common distinct modeling type (excluding ensemble which will be discussed below) was the category of linear models, mixed models, or BLUP type models that were used by 52% of respondents. These linear statistical methods are the basis of much modern breeding, making their prominence in the competition unsurprising. As with classical machine learning, these methods were often combined with other strategies to create a final prediction. Both environment inclusive and agnostic methods were applied within this category. The winning team’s methods were based primarily on this category of modeling frameworks.

Deep Learning and/or Neural Network based methods were the third most common single modeling strategy at 33%. One particularly unique approach in this category came from the team “Niche Squad” which used Vision Transformers (Dosovitskiy et al. 2021), a method from image classification, as the bases for their neural network models. While the models did not perform well in the competition, the team identified many improvements that could be made to the model with greater time and effort than was possible during the short competition window.

Two teams in the competition used other strategies that did not fit into the categories provided. “SmAL”, used a deterministic model in combination with deep learning approaches to achieve a seventh-place finish in the competition, whereas the other team (ranked 20^th^) did not share their method. Ensemble approaches, especially simple ones based on averaging model predictions together, were particularly prevalent in the competition with 52% of respondents opting to combine multiple types of models in this way. Some of these uses were as simple as averaging the results of replicated runs of the same non-deterministic model, while other used ensembling to combine the results from different model types that could be hypothesized to rely more strongly on different aspects of the G2F dataset.

Another important decision each team was required to make was which of the data types provided by the competition, and any other public datasets, they would use in their model and how those data would be represented (Table 4). Genetics was the number one factor used by 93% of respondent teams. There were only two teams that did not use genetics in their models, although another team initially excluded genetics from their model but then added it in and found accuracy improvements (See File S2). Surprisingly, one of the teams that explicitly excluded genetic factors ranked in second place in the competition. Importantly though, the environmental covariates provided by the competition and used by this team were generated by a method that employed some limited use of genetic information (Lopez-Cruz et al. 2023). Additionally, maize hybrids are typically adapted to different climates across the US, often performing poorly outside of those locations. For this reason, the G2F GxE project applies a stratified approach where not all hybrids are grown in every location and the hybrids grown in any given location are oversampled for those that are adapted to that location’s climate. This likely amounts to some genetics being represented in factors that would otherwise be considered strictly environmental. Additionally, past work on the G2F GxE datasets has indicated substantially more variance among environments than among hybrids, due to the large range of environmental conditions: much larger than a typical hybrid development trial (Lopez-Cruz et al. 2023; Rogers et al. 2021; Rogers and Holland 2022; Washburn et al. 2021). Another important factor is that the competition was based on a very difficult prediction scenario where many of the hybrids in the testing set did not exist in the training set, and even the parents of hybrids in the testing set were largely different from those in the training set. Because of these differences, even the models that explicitly included genetic factors may have struggled to make significant accuracy improvements using them. Regardless of the reasons, this second-place model demonstrates that it is possible to make reasonable predictions based on limited genetic input in the context of the G2F GxE project, even if these predictions would probably not be particularly useful in a breeding context. It is worth noting as described in the introduction that many important research and application questions can and have been addressed with E/M-centric modeling approaches.

**Table 4.** Factors included in the models by team.

|  | Factors Included in Model |  |  |  |  |  |  |  |
| --- | --- | --- | --- | --- | --- | --- | --- | --- |
| Team Name | Rank | Genetics | Weather | Soil | Environment Covariates | Field Management | Experimental Design | Other Factors |
| CLAC | 1 | X | X | X | X | X |  |  |
| igorkf | 2 |  | X | X | X | X |  |  |
| phenomaize | 3 | X |  |  |  |  | X |  |
| UCD_MegaLMM | 4 | X |  |  |  |  | X |  |
| CGM | 5 | X |  |  |  |  |  | X |
| Purdue | 6 | X | X | X |  |  | X |  |
| SmAL | 7 | X | X | X | X | X |  |  |
| ML_APT | 8 | X | X | X | X | X | X |  |
| MPB_Group | 9 | X | X | X |  | X |  | X |
| AlBreeding | 10 | X | X | X | X |  |  |  |
| AgroStat | 11 | X | X |  |  |  |  |  |
| arulrich | 12 | X | X | X |  |  |  |  |
| DataJanitors | 13 | X |  |  | X | X | X |  |
| CropsAreCool | 14 |  | X | X | X | X | X |  |
| AllModelsAreWrong | 15 | X | X | X | X |  |  | X |
| agAdaptAR | 16 | X | X | X | X |  |  | X |
| DeepCropVision | 18 | X | X | X | X | X |  | X |
| supermanwasd | 19 | X | X | X | X |  |  |  |
| BioSense | 20 | X | X | X | X | X |  |  |
| Kernel of Truth | 21 | X |  |  |  | X |  |  |
| AIMaize | 24 | X | X | X | X |  |  |  |
| EnBiSys | 25 | X | X | X | X | X |  |  |
| Genetwister | 26 | X |  |  |  |  | X |  |
| gartybois | 27 | X | X | X | X | X | X |  |
| Niche Squad | 28 | X |  | X | X |  |  |  |
| uwaBioinfo | 29 | X | X | X |  |  |  |  |
| TinyAfrica | 30 | X | X | X | X | X |  | X |
| Percentage of Teams Using Factor |  | 93% | 74% | 74% | 63% | 48% | 30% | 22% |

Weather and soil data were the next most highly used factors at 74% each. Along with the environmental covariates, used by 63% of respondents, these constituted the strictly environmental factors (if human controlled management is considered separately from non-human-controlled environment) provided by the competition. Field management (tillage, irrigation, etc.) and experimental design factors, which include details like field replicates and spatial arrangements in the field, were used by 48% and 30% of respondents, respectively. Additionally, 22% of the teams included other factors such as external historical datasets in their models.

### Winning team’s strategy and results

The winning team’s strategy involved: 1) Identifying and defining the target population of environments (TPE) and target population of genotypes (TPG) to better understand the prediction challenge. 2) Considering the implications of the evaluation metric used in the competition (i.e., RMSE favors absolute differences whereas Pearson r is a correlation and favors rank differences). 3) Focusing their efforts on modeling location means and genotypic performance separately. The team reasoned that getting the environmental means within a similar scale to the observed values might pay off more than predicting the ranks of genotypes within each environment correctly. Moreover, any modeling artifacts, such as “shrinkage” of coefficients, would be more detrimental to RMSE scores than it might be to Pearson r or other rank-based metrics.

Figure 5 displays the spatial distribution of environments, as year-location combinations, and the prediction targets. All locations in the testing set had already been observed in the previous years of data utilized to calibrate the model (training set), indicating that a “location” term could be directly used as a predictor in the model for location means, as well as an informative way to model GXE interactions. Thus, the unobserved environmental means were primarily informed by the average location performance from previous years, complemented by the available information in the provided metadata (e.g., treatment, previous crop, etc.) treated as covariates, and environmental covariates as captured by a random forest model.

**Figure 5.**
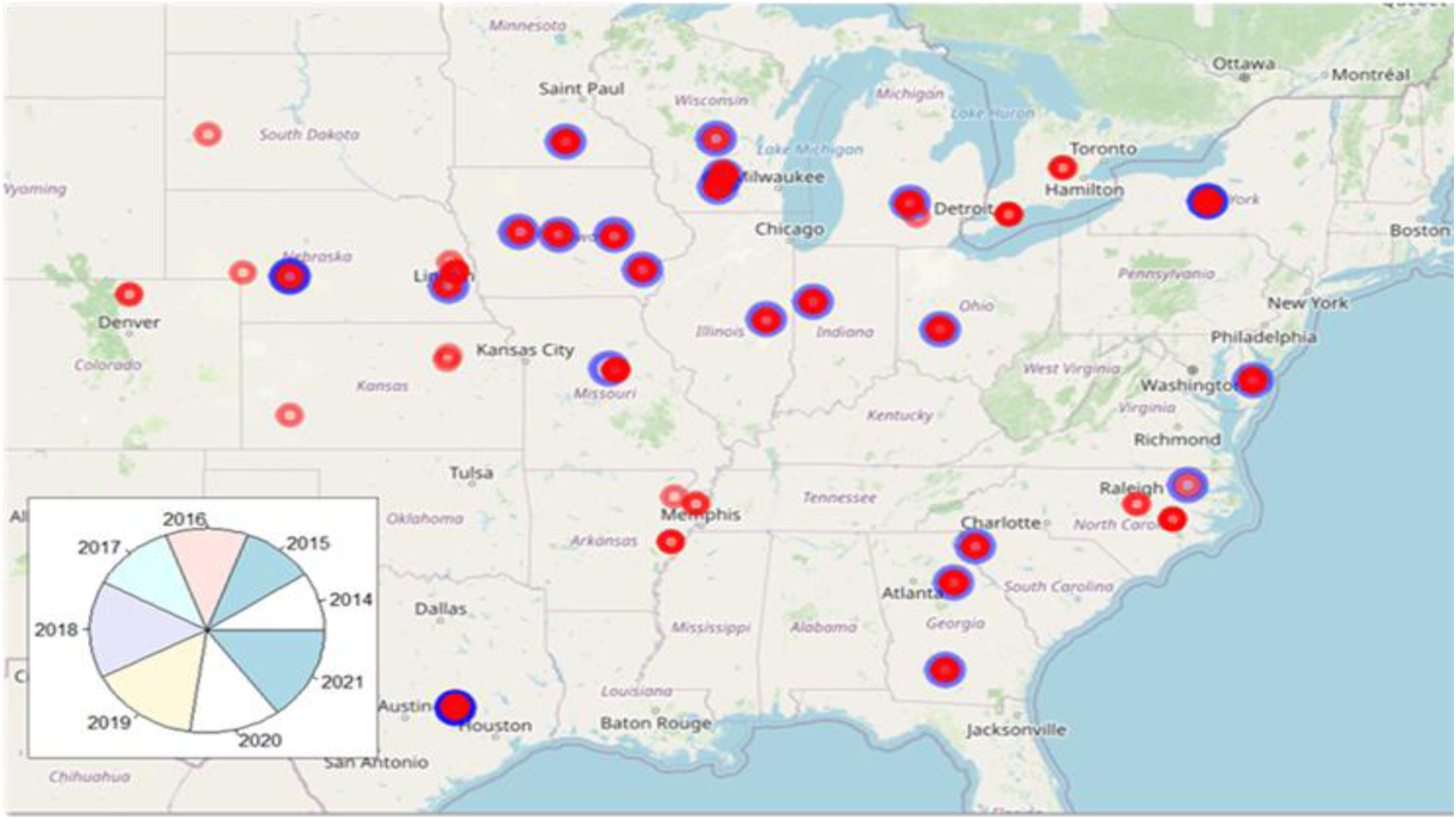
Target population of environments: Distribution of trial data (red dots) and prediction targets (blue circles). The pie chart indicates the distribution of data points per year.

Regarding the target population of genotypes, Figure 6 indicates that the prediction target is contained within the genotypic space of the training data based on the two first principal components, although it is not necessarily well represented. Genomic information was provided at the hybrid level, and the clouds seem to indicate substantial contribution of testers to the population structure. Relationship information was utilized by the winning team to infer the determinist accuracy between every pair of locations, specifically between observed and target environments. Together, deterministic accuracy estimates from genomic information harnessing the TPG, and the spatial location, capturing the TPE were used to assign the contribution of each training environment to the prediction of each testing environment. Since the shrinkage of genomic values was likely to have a detrimental impact on the evaluation metric, predicted genomic values (from marker information) were normalized and rescaled. This was done using a standard deviation value the team predicted for each environment based on a model analogous to the location mean model.

**Figure 6.**
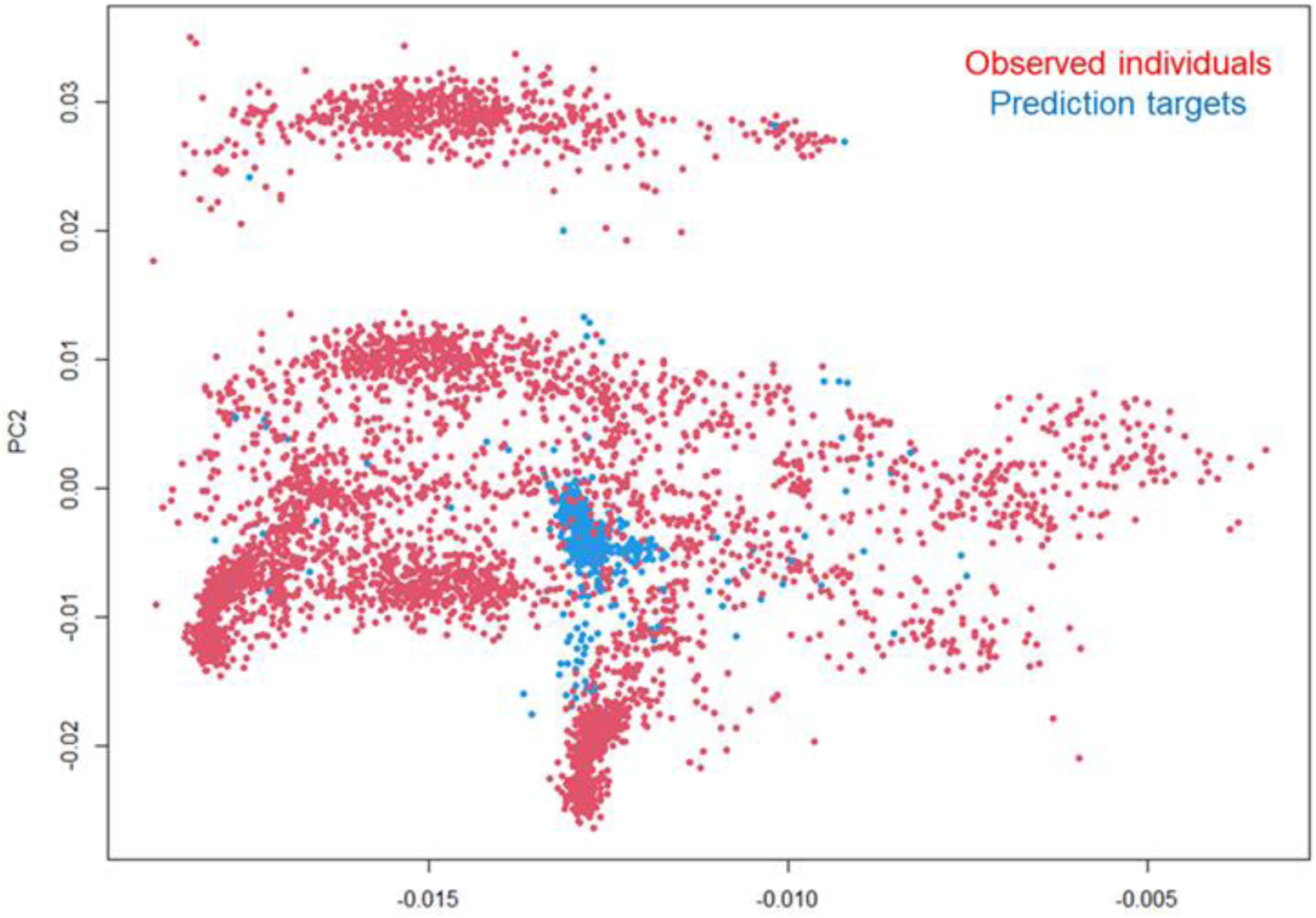
Target population of genotypes: First and second principal components of the genomic relationship matrix with observed individuals (red dots) and prediction targets (blue dots).

Each of the winning team’s submissions is listed in table 5. The modifications made between each submission were based on the feedback (RMSE scores) from the previous submission and conjectures made by the team. To gain degrees of freedom and predictive power, the data concerning previous crops were aggregated into four levels: wheat, legume, corn or other. Treatments were also aggregated into standard, dry and late. Irrigation information (yes/no) was removed from treatments, and it was attributed to an additional covariate.

**Table 5:** Winning team’s modeling strategy by submission.

| Submission | RMSE | r | Model Description |
| --- | --- | --- | --- |
| S1 | 2.454 | 0.263 | GBLUP with minor QC and some environmental covariates |
| S2 | 2.521 | 0.260 | GBLUP with QC on GEBVs and location means; no environmental covariates |
| S3 | 2.353 | 0.266 | Ensemble: Average of methods used in S1 and S4 |
| S4 | 2.410 | 0.267 | GBLUP with QC on GEBVs; No location means QC; No environmental covariates |
| S5 | **2.329** | **0.357** | Ensemble: Average of S1 and univariate GBLUP |

The team’s first submission was based on their best guess and reasoned strategies from prior experience: a multivariate model with minimal data QC. With the hope of improving this model for their second submission, members of the team with breeding knowledge reviewed the data and applied more stringent quality control (QC). This involved dropping outliers using a 3-standard deviation rule, removing locations with treatments tagged as ‘disease trial’, and removing experimental units with stand counts below 20. However, this second submission with greater QC performed worse than the first one, possibly because the quality of the testing set was comparable to the training data, causing the data QC to make the dataset less representative of the prediction target. At this point the team was uncertain how to proceed but determined to submit their first model averaged with a slightly simpler model without any environmental covariates (ECs). This resulted in a better RMSE than either of the first two submissions. To determine if the simpler model was better by itself, they submitted it alone as their fourth attempt. The poorer result indicated it was not. For their final submission, the team used their original model (from submission one) but averaged it with an even simpler univariate GBLUP than submission four, resulting in the winning score. Although the winning team’s strategy was in part focused on predicting environmental means, their final winning submission actually resulted in a large increase in Pearson r over previous submissions (Figure 7) indicating that better within environment (genetic) prediction was a component of what boosted them to the winning position. After further post-competition study they concluded that because the data was very structured, most of the predictive ability within-environment was probably coming from the univariate GBLUP rather than the unstructured/multivariate GxE model.

**Figure 7.**
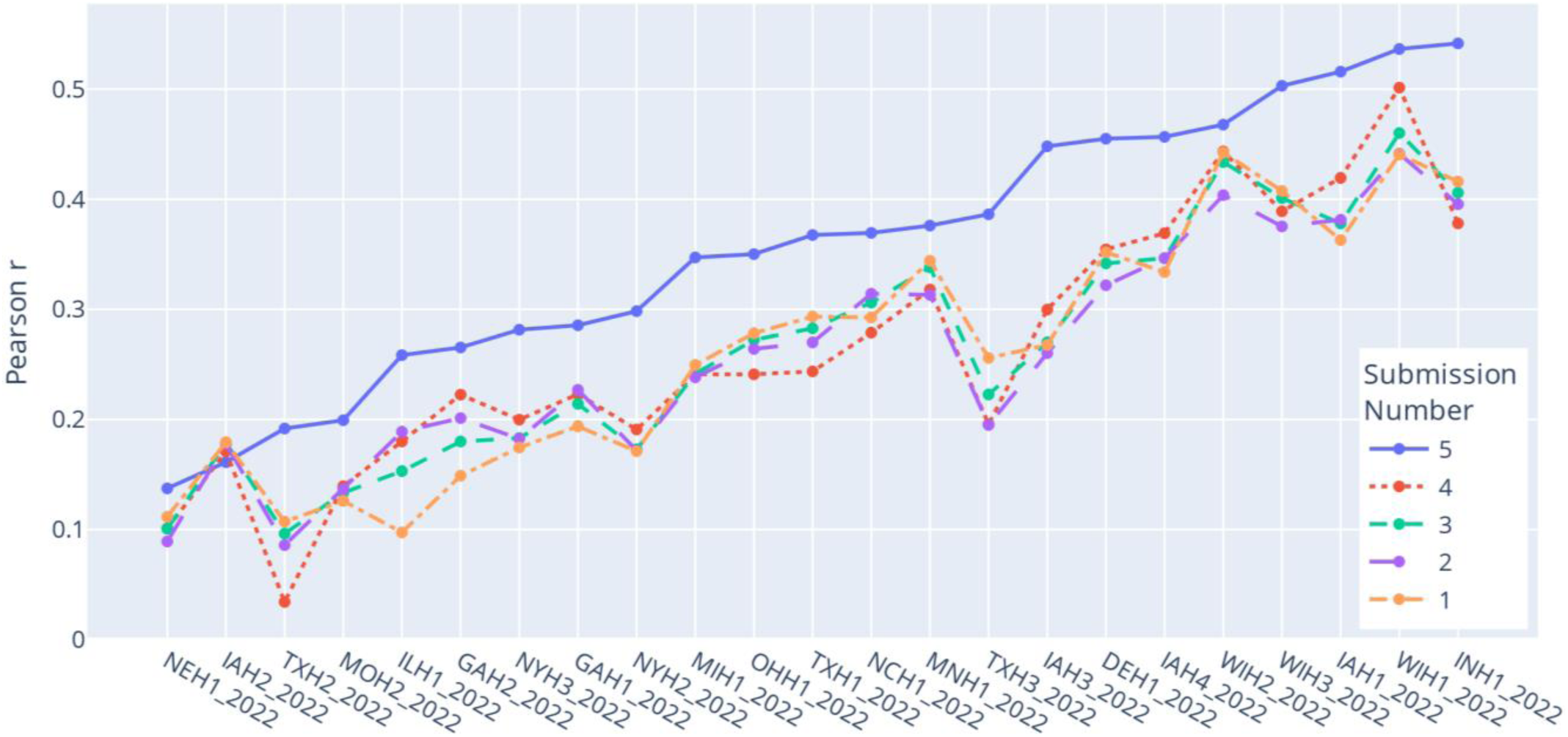
Pearson r values from the winning team by location and submission.

### Conclusions

A large diversity of model types, strategies, and data inputs were used in the competition and, perhaps surprisingly, many of these made it into the top ten list. This demonstrates that no single modeling strategy is greatly superior to the others, at least in the context of this prediction problem. Additionally, ensemble approaches, even those as simple as averaging, combining multiple models with different strengths and weakness often outperform single models. The winning team’s strategy of attempting to model environmental means separately from genetics appears to have been effective with the simple univariate GBLUP from the final model improving both absolute (RMSE) and relative/rank-based (Pearson r) scores within most environments.

The level of international interest and participation in the competition demonstrates that there is broad excitement about crop yield prediction from many disciplines and carrier stages. Crop yield prediction comes with both significant challenges and enormous potential benefits to society. Although the bulk of registrants came from plant science, genetics, or breeding backgrounds, participants with little or no background in these areas also participated. Activities like this competition have the potential to bring new ideas and ways of thinking from disciplines like computer science and engineering into the genetics and plant science communities, enhancing our abilities to solve critical, and technically challenging, problems.

## Data Availability Statement

All data used in this manuscript are publicly available at https://doi.org/10.25739/tq5e-ak26. Code from all teams is publicly available as follows:

**AgAdaptAR**: https://github.com/EcoEvoInfo/maize-gxe-prediction-challenge-2023,

**AIMaize**: https://github.com/ksegaba/Genomes2Field_Competition,

**All Models are Wrong**: https://zenodo.org/record/7830071,

**arulrich**: https://github.com/mwylerCH/GxEcompetition,

**CLAC**: https://github.com/alenxav/Lectures/tree/master/MGC_2023,

**DataJanitors**: https://github.com/qchen33/g2fcompetition2022,

**DeepCropVision**: https://github.com/Ved-Piyush/DeepCropVision_maizegxeprediction2022,

**EnBiSys**: https://github.com/dperondi/maizegxeprediction2022,

**gartybois**: https://github.com/Thyra/g2f-maize-challenge-2022,

**Kernel of Truth**: https://github.com/robertkhu/maize_gxe,

**ML_APT**: https://forgemia.inra.fr/ml_apt/g2f_challenge,

**MPB_Group**: https://zenodo.org/records/12721443,

**Niche Squad**: https://github.com/Niche-Squad/gsformer,

**Phenomaize**: https://forgemia.inra.fr/renaud.rincent/genome2fields,

**SmAL**: https://github.com/SmartAgriLabs/G2F-competition,

**UCD_MegaLMM**: https://github.com/ucdavis/UCD_MegaLMM,

**uwaBioinfo**: https://github.com/eyesoftruth/G2F-competition2022.

## Acknowledgments

We sincerely thank the field managers, crop research coordinators, staff, graduate students, student interns, and data collectors for their efforts in the GxE project. Team phenomaize is grateful to the INRAE MIGALE bioinformatics facility (MIGALE, INRAE, 2020. Migale bioinformatics Facility, doi:10.15454/1.5572390655343293E12) for providing computing resources.

